# *In situ* genome editing method suitable for routine generation of germline modified animal models

**DOI:** 10.1101/172718

**Authors:** Masato Ohtsuka, Hiromi Miura, Naomi Arifin, Shingo Nakamura, Kenta Wada, Channabasavaiah B. Gurumurthy, Masahiro Sato

## Abstract

Animal genome engineering experimental procedures involve three major steps: isolation of zygotes from pregnant females; microinjection of zygotes, and; transfer of injected zygotes into recipient females, that have been practiced for over three decades. The laboratory set ups intending to performing these procedures require to have sophisticated equipment as well as highly skilled technical personnel. Because of these reasons, animal transgenesis experiments are typically performed at centralized core facilities in most research organizations. We recently showed that all three steps, of animal transgensis, can be bypassed using a method termed GONAD (<u>G</u>enome-editing via <u>O</u>viductal <u>N</u>ucleic <u>A</u>cids <u>D</u>elivery), by directly electroporating genome editing components into zygotes *in situ*. Although our first report demonstrated the genome-editing capability, its efficiency was lower than the standard methods using microinjection. Here we investigated critical parameters of GONAD to make it suitable for creating animal models of large genomic deletions, single nucleotide corrections and long sequence insertions. The efficiency of genome editing in the improved GONAD (*i*-GONAD) method reached to the levels comparable to traditional microinjection methods. The streamlined parameters, and the simplified experimental steps, in the *i*-GONAD method makes it suitable for routine genome editing applications performed both at centralized facilities as well as at the laboratories that lack highly skilled personnel and the sophisticated equipment.

## Introduction

Recent advents in the development of genome editing such as CRISPR/Cas9 enables production of knock-out mice easily and rapidly ^1–3^. The experimental procedures of CRISPR genome engineering include three steps: isolation of zygotes from super-ovulated females that were previously mated with males, zygote microinjection of genome editing components, and transfer of microinjected zygotes into the oviducts of pseudopregnant females ^1,2^. These steps require: (i) very high level of technical expertise, proficiency and skills among the technicians who perform these procedures; and (ii) expensive apparatus such as micromanipulators. Because of the complex nature of these experimental steps, animal genome engineering experiments are difficult to perform in many laboratories, and are typically performed in centralized core laboratories, in most research organizations, where trained personnel offer genome engineering services on a day-to-day basis. Developing of methods that can avoid such complex steps can make the animal genome engineering technologies to be performed at many laboratories, not just core laboratories. Some groups have investigated whether *in vitro* electroporation of zygotes can be used as an alternative to microinjection, and they have been successful in producing genome-edited fetuses and pups using this approach ^4–9^. Electroporation of zygotes overcomes the microinjection step (the middle of the three steps), but the strategy still requires the other two steps: isolation of zygotes for *ex vivo* handling and their transfer back into females. We recently demonstrated that all the three steps can be bypassed by performing electroporation of zygotes *in situ*.

To further simplify germline genome editing, we recently developed a method called <u>G</u>enomeediting via <u>O</u>viductal <u>N</u>ucleic <u>A</u>cids <u>D</u>elivery (GONAD), which does not require isolation of zygotes, nor their *ex vivo* handling for microinjection and subsequent transfer to recipient females (10). GONAD is performed on E1.5 day (2-cell stage) pregnant females. The ovaries and oviducts are surgically exposed through an incision at the dorsolateral portion, and genome editing reagents are instilled into oviductal lumen using a glass capillary pipette. Immediately after solution instillation, the entire oviduct is subjected to *in vivo* electroporation using tweezer-type electrodes. After electroporation, the ovary-oviduct parts are returned to their original position and incision closed by suturing. The *in situ* genome-edited embryos will develop to full term and offspring are genotyped for targeted mutation. We demonstrated, through a proof of principle report, that it is possible to create *indel* mutations at the target loci in some of the fetal offspring with 28% efficiency (7/25) (10). At the time of developing this strategy we realized that the method can be significantly improved, by systematically testing various parameters, to enable the method to achieve precise genome editing. The improvements that needed to be achieved or tested were whether: (1) small point mutation knock-in and large cassette knock-ins can be created (not just creating *indels*); (2) induced mutations in the founder (G0) pups can be transmitted to the next generation; (3) mosaicism, which typically occurs if genome editing happens at 2-cell stage and beyond, can be reduced; (4) GONAD can be performed using other commercially available electroporators (the model we used in our initial experiment is no longer available and has not been made by the manufacturer for over a decade); 5) females subjected to the GONAD procedure retain their reproductive function; and (6) GONAD can be performed using Cpf1, the second-most commonly used CRISPR family nuclease.

In this study, we made major modifications of the GONAD to improve the above mentioned points. We termed the modified GOAND as *i*mproved GONAD (*i*-GONAD). The *i*-GONAD method offers much higher genome editing efficiencies. We demonstrate that *i*-GONAD approach can be used for creating germline modified G1 offspring with genetic changes such as large deletions and knock-ins. Furthermore, we demonstrate that the *i*-GONAD method is robust because it works with commonly used electroporators. These features makes *i*-GONAD to a easily adaptable at many laboratories and can be performed by many researchers including beginners or students who do not typically possess special skills to operate a micromanipulator.

## Results

### 1) GONAD on Day 0.7

In our first report of GONAD method development, experiments were performed at ~1.5 to 1.7 day post mating. This stage of pregnancy corresponds to 2-cell embryos. Delivery of genome editing reagents at this stage can result in higher frequency of mosaic embryos and/or fetuses ^10^. The ideal time of delivery would be the one that corresponds to 1-cell stage zygotes because it can reduce mosaicism events. In order to investigate the earliest time of delivery, we tested GONAD at two separate time points; Day 0.4 and 0.7. We instilled 1.0~1.5 µl of a solution containing eGFP mRNA (1 µg/µl) and trypan blue into oviductal lumens (as shown in **Fig. 1A**, schematically) and subsequently performed *in vivo* electroporation using a BTX T-820 electroporator with the conditions we previously described ^10^. Two days after gene delivery, eight cell-stage embryos were isolated from the treated females and observed for eGFP fluorescence. We did not observe appreciable eGFP signals in any of the embryos isolated from 0.4 day time point, whereas uniform fluorescence was readily observed in 13 of 31 embryos isolated from 0.7 day time point females (**Fig. 1B**). One of the reasons for lack of success at Day 0.4 could be that the zygotes are tightly surrounded by cumulus cells at this stage (**Fig. 1C**, right), and this may hamper efficient electrophoretic delivery of nucleic acids to zygotes. On the other hand, most zygotes at Day 0.7 would be devoid of thick cumulus cells surrounding them (**Fig. 1C**, right) to permit the injection mix fluids to readily reach zygotes. These results suggested that 0.7 day (~16 hours) post-mating would be a most suitable time for GONAD procedure.

**Fig. 1.**
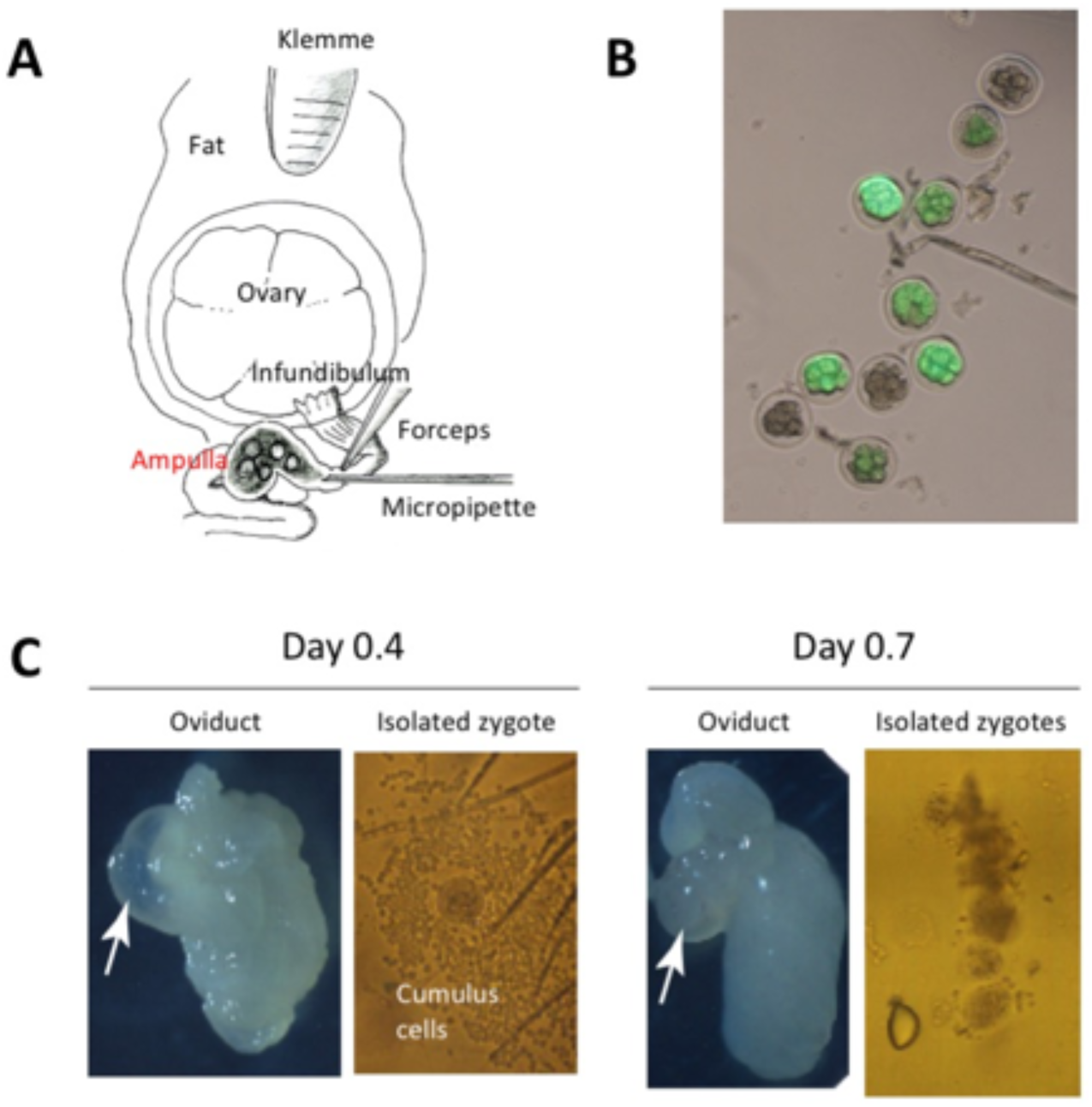
Evaluation of earlier time points for performing GONAD. **A.** Diagrammatic illustration showing the anatomical structures of ovary, oviduct and the surgical equipment used for GONAD procedure. A small amount of solution is injected by direct insertion of a glass micropipette through oviductal wall located at the region between the ampulla and infundibulum. Immediately after injection, *in vivo* electroporation is performed on the entire oviduct. **B.** Detection of eGFP fluorescence in 8-cell to morula embryos after delivery of eGFP mRNA via GONAD procedure. The eGFP fluorescence in morulae, isolated two days post GONAD procedure performed on naturally mated ICR female at 0.7 day of pregnancy. **C.** Oviducts and zygotes dissected on Days 0.4 (left panel) and 0.7 (right panel). Note that the oviduct dissected on Day 0.4 exhibits swelling of the ampulla (arrow). The zygotes isolated from the Day 0.4 ampulla are usually surrounded by thick layer of cumulus cells. These cells may hamper efficient uptake of exogenous nucleic acids/proteins when it is instilled intra-oviductally and subsequently electroporated. The oviduct dissected on Day 0.7 exhibits shrinkage of the ampulla (arrow), and zygotes isolated from the Day 0.7 ampulla have few numbers of cumulus cells, which will less likely hamper the uptake of exogenous nucleic acids/proteins upon electroporation.

### 2) Production of *Foxe3* knock-out lines by the modified GONAD

Next, we tested if the GONAD method can be used for creating gene disrupted animal models. We chose *Foxe3* locus for gene targeting (**Fig. 2A**), inactivation of which is known to cause abnormal development of the eye and cataract in mice ^12,13^. GONAD was performed in 0.7 day pregnant ICR females using Cas9 mRNA and a single-guide (sg)RNA against the *Foxe3* gene. The electroporation conditions used were 50 V in voltage, 5 msec of wave length and 8 pulses. Five of the eight GONAD-treated females delivered pups. Sequencing analysis of genomic DNA isolated from ears of G0 offspring demonstrated that 11 of 36 G0 pups (31%) had mutated allele in the target locus (**Figs. 2B** and **2C**; **Supplemental Table 1**). Notably, 55% of these (6 of the 11) pups exhibited mosaicism alleles, and mosaicism was not detected among the remaining five pups (e.g. pups G0-#26 and -#32 in **Fig. 2B**). This level of mosaicism is much lower than that observed in samples derived from GONAD performed on Day 1.4 ^10^. Intercrossing of G0 founders resulted in G1 offspring, in which some exhibited the expected cataract phenotype (**Figs. 2D** and **3E**). Sequencing of genomic DNA isolated from G1 mice showing with cataracts revealed germ-line transmission of mutated alleles (**Figs. 2C-2E**). Taken together, these results indicate that the GONAD method can be used for creating gene disrupted mouse models.

**Fig. 2.**
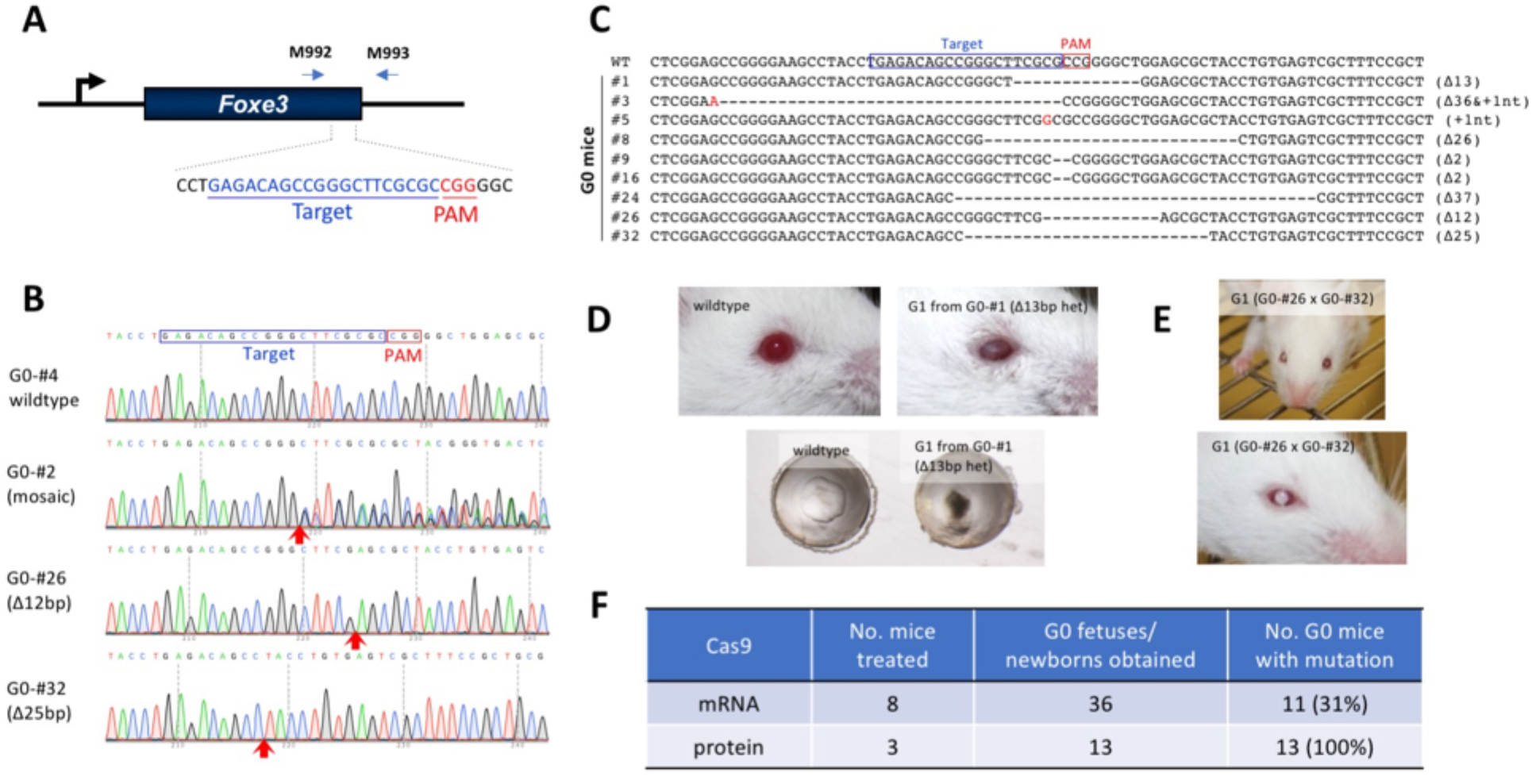
Creating gene inactivated animal models using the GONAD method. **A.** Schematic of the targeting strategy to inactivate *Foxe3* gene and the primer set used for genotyping. **B.** Direct sequencing results of PCR products amplified from the founder (GO) mice with the primer set shown in **A.** The red arrows below the electropherogram show the region with *indel* mutations. **C.** Mutated *Foxe3* alleles possessed in the GO mice. The changes in the nucleotide sequence are shown in red and the type of changes (insertions: +Xnt, or deletions: Δ) are indicated on the right-side of the sequences. **D** and **E.** Cataract phenotypes in the G1 mice. **F.** Efficiencies of *Foxe3* gene editing. CRISPR components used were either Cas9 mRNA/sgRNA or Cas9 protein/crRNA/tracrRNA (see **Supplemental Table S1** for detail).

### 3) Higher genome editing efficiency using crispr RNA + tracr RNA + Cas9 protein (ctRNP) complexes

It is now becoming increasingly clear that the use of Cas9 protein, instead of Cas9 mRNA ^7,14,15^, and, crRNA + tracrRNA (two part guide RNA), instead of single guide RNA (sgRNA), yield higher genome-editing efficiencies ^14^. We thus examined the combinatorial use of these components (crRNA + tracrRNA + Cas9 proetin: ctRNP) for GONAD-mediated genome editing. We used the *Foxe3* gene, as an example, using the same guide sequence as in the previous experiment except that, annealed crRNA + tracrRNA were used in place of sgRNA and Cas9 protein was used in place of Cas9 mRNA. A mixture of ctRNP complexes was instilled into the oviductal lumen of three pregnant ICR females at Day 0.7, and then electroporated *in vivo* using the same conditions as before. The embryos were isolated at E17.5. All the three females contained fetuses and a total of 13 G0 fetuses were obtained from them. Surprisingly, all the 13 fetuses (13/13, 100%) had *indel* mutations within the target sequence of the *Foxe3* gene (**Fig. 2F**; **Supplemental Table 1**). In direct comparison, when Cas9 mRNA was used, genome editing efficiency was only 31%. These results clearly suggest that combinational use of crRNA, tracrRNAs and Cas9 protein (ctRNP) offers the highest efficiency of genome editing in GONAD. We termed this RNP-based GONAD as *i*mproved GONAD (*i*-GONAD).

### 4) Creation of gene corrected animal models using *i*-GONAD

Next, we tested if *i*-GONAD method could be used for making small genetic changes to rescue a phenotype. We chose G308C mutation in the *tyrosinase* (*Tyr*) gene as an example for this experiment. This mutation changes cysteine to serine at amino acid 103 of the tyrosinase protein, leading to its defective activity, and the phenotype of this genetic change can be assessed by eye pigmentation (clear or dark) or coat color (albino or agouti) in ICR strain mice ^16^. We designed a gRNA for a region spanning the point mutation and constructed ssODN that corresponds to the wild-type sequence of *Tyr* (**Fig. 3A**).

**Fig. 3.**
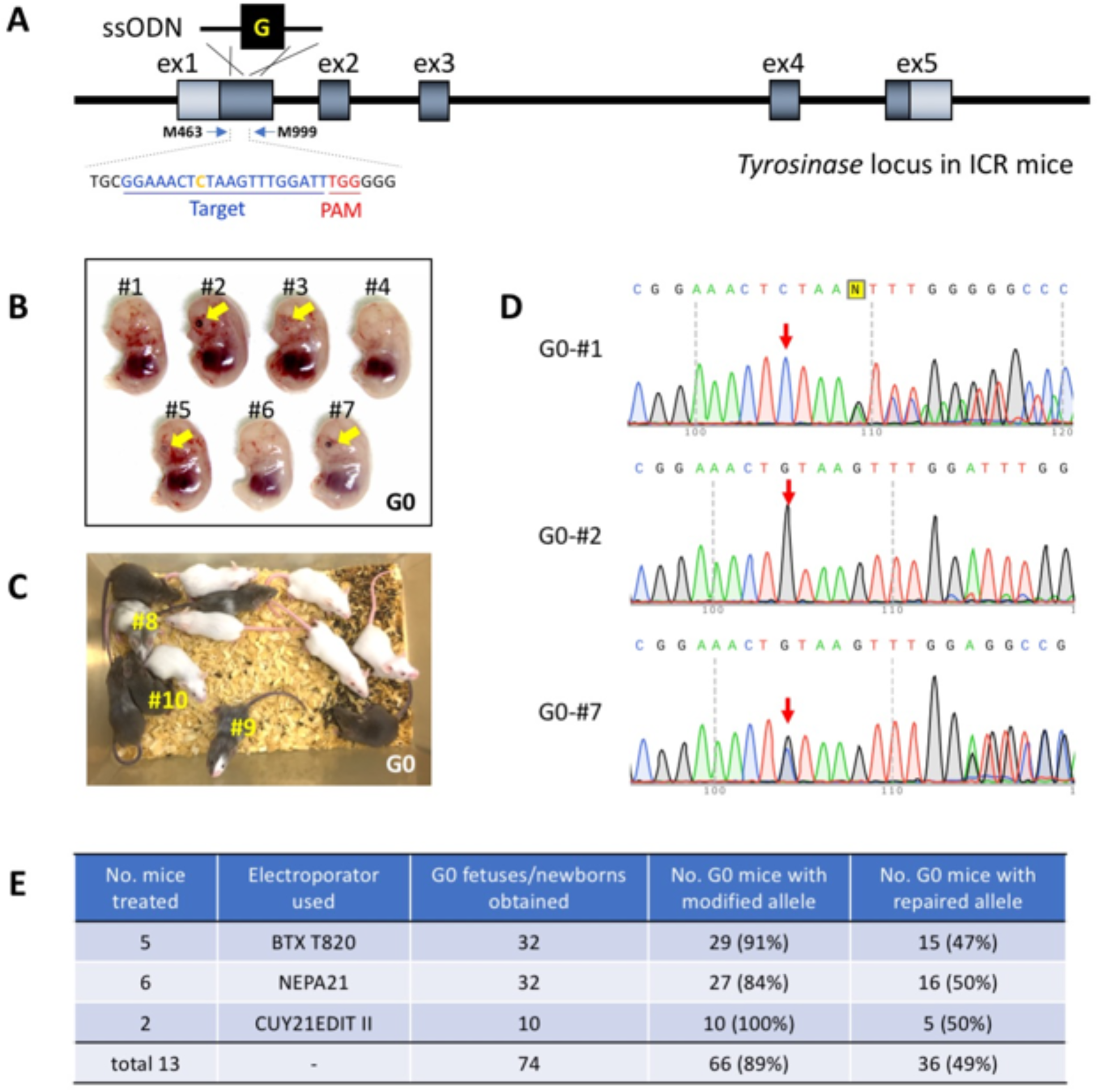
Creating small genetic change animal models using the GONAD method. Restoration of *Tyr* gene of albino ICR mice by ssODN-based knock-in with the GONAD method. **A.** A schematic to show rescue of *Tyr* gene mutation. The target region containing the guide sequence and the genotyping primer binding sites are shown. **B.** Representative E14.5 litter showing *Tyr* rescued GO fetuses. The pigmented eyes of the fetuses are indicated by yellow arrows. **C.** Representative *Tyr* rescued GO mouse litters obtained from #5 female mouse in **Supplemental Table S2**. GO mice indicated in # numbers (shown in yellow) were used for germline transmission analysis (see details in **Supplemental Fig. S2**). **D.** Direct sequencing results of PCR products amplified from the GO fetuses (in B). The position for mutated/corrected nucleotides are indicated by red arrows. **E.** Efficiencies of *Tyr* gene editing. Three kinds of electroporator were used in the experiment (see **Supplemental Table S2** for details).

*i*-GONAD was performed in five pregnant ICR females. A total of 32 offspring from these females was harvested at different stages of gestation (from E14.5 to E19.5), or at post-natal stage. Fifteen (47%) of these samples exhibited the expected phenotype of eye pigmentation (in fetuses) or coat color (in newborn pups) (**Figs. 3B** and **3C**). Sequence analysis demonstrated that at least one allele had the corrected sequence (G to C) at the target site in all of the offspring with a pigmentation phenotype (**Fig. 3D**; **Supplemental Table 2**).

The experiments were performed using the BTX T820 electroporator, a model that, as noted, is no longer manufactured. We therefore tested two newer electroporators, NEPA21 (Nepa Gene Co) and CUY21EDIT II (BEX Co). Six females were subjected to *i*-GONAD using NEPA21, yielding 32 offspring of which 16 (50%) showed the expected genetic change, including the phenotypic change of eye pigmentation or coat color (**Fig. 3E**; **Supplemental Table 2**). Two females were subjected to *i*-GONAD using CUY21EDIT II, and fetuses were analyzed at E14.5. Of the 10 fetuses obtained, 5 (50%) showed the expected genetic change as well as phenotypic change (eye pigmentation or coat color) (**Fig. 3E**; **Supplemental Table 2**).

Highly consistent genome editing efficiencies (47%, 50% and 50%) obtained from three different electroporators (**Supplemental Fig. 1)** demonstrates the robustness and reproducibility of the *i*-GONAD method. These results suggest that *i*-GONAD can be used for high efficiency gene correction via co-delivery of ssODN repair donor.

Eighty-three % (30/36) of the offspring with the pigmentation phenotype were found in these three experiments to have *indel* mutations around the target region in their second alleles (**Fig. 3D**). Of note, 79％ (30/38) of the unpigmented offspring, that were thought to be not rescued had *indel* mutations. Considering gene correction and *indels* together, a total of 89% (66/74) of G0 offspring were genome-edited (**Fig. 3E**). These results again support that *i*-GONAD can yield very high efficiencies of genome editing.

The representative three founder mice containing repaired allele (G0-#8: low contribution mosaic, G0-#9: 60% mosaic, G0-#10: 100% mosaic [based on the coat color]; **Fig. 3C**) were bred with ICR mice to assess germline transmission of repaired allele. Although we could not obtain rescued progeny from G0-#8 (0/43), pups exhibiting agouti coat color were obtained from founders G0-#9 and -#10 (#9 [8/18] and #10 [30/30]) (**Supplemental Figs. 2A** and **2B**). From the germline transmission rate of G0-#10, it should have rescued allele as homozygotes, as indicated by the sequencing result (**Supplemental Fig. 2C**).

To directly compare the efficiencies of *i*-GONAD with standard microinjection-based approach, we performed *Tyr* gene correction through microinjection of ctRNP and ssODN donor into zygotes isolated from females. We collected oocytes from 8 superovulated ICR females that were subjected to *in vitro* fertilization (IVF) to generate zygotes for microinjection. Sixty-four zygotes were injected with CRISPR reagents and cultured, of which 50 (78%) advanced to the 2-cell stage, and these were then transferred to two recipient females. The females were euthanized at E14.5 for embryo harvesting: one female held three fetuses and the second female did not hold any. Only one of the three fetuses (33%) had pigmented eyes. Sequencing analysis showed that the fetus with pigmented eyes had the corrected sequence at the target site (**Supplemental Table 3**), but the other two fetuses, with non-pigmented eyes, had neither corrected sequence nor the *indel* mutations around the target site. These data, of direct comparison of point mutation genome editing using *i*-GONAD and standard microinjection-based experiments, clearly demonstrates the potential merits and superiority of *i*-GONAD approach at least in for the ICR strain. Notably, the standard microinjection-based approach needed about three times more animals for the experiment (8 females as egg donors + 1 male as sperm donor for IVF + 3 pseudopregnant females + 3 vasectamized males = 15 mice), whereas *i*-GONAD used only 6 mice (3 embryo donors mated with 3 stud males = 6), yet created correctly genome edited models at a higher efficiency than the microinjection approach.

### 5) Generation of large genomic deletion animal models using *i*-GONAD

We next examined whether the *i*-GONAD method can be used for production of mice with large genomic deletions. We targeted a retrotransposon sequence present in the 1st intron of *agouti* locus in C57BL/6 strain mice ^17^. A large deletion, spanning 16.2 kb, containing the retrotransposon sequence, was intended to be deleted by making two cleavages flanking the retrotransposon insertion site (**Fig. 4A**). The deletion of this region would result in a coat color change from black to agouti in C57BL/6 mice (**Fig. 4B**). To aid precise joining of the genomic ends, we provided an ssODN donor containing short homology sequences for the two cleaved ends. The ssODN also included an *Eco*RI site for using it in an RFLP assay. We instilled a solution containing Cas9 protein, two crRNA/tracrRNAs and ssODN into the oviductal lumen of eight E0.7 pregnant C57BL/6 females and performed *in vivo* electroporation. Of these, only four females remained pregnant, from which six G0 pups were recovered through Caesarean section. Genotyping the pups revealed that 3 (50%) of them exhibited large deletion in their *agouti* locus (G0-#3 and -#5) and/or agouti coat color (G0-#4 and -#5) (**Fig. 4B** to **4D**; **Supplemental Table 4**). Notably, one pup (G0-#3; 1/6 [17%]) had correctly inserted ssODN at the target site, while the another (G0-#5) showing agouti coat color did not have the insertion (**Fig. 4D** and **4E**). The PCR did not yield an amplicon from the G0-#4 agouti pup, suggesting that one or both of the primer binding sites may have been deleted. Interestingly, the pup G0-#6, one of the pups that neither showed large deletion at the target region nor agouti coat color, had *indel* mutations at both the cleavage sites (**Fig. 4F**; **Supplemental Table 4**). The percentage of individual pups that had been genome-edited (large deletions and short *indels* together) was 67% (4/6). These data suggest that *i*-GONAD can create large genomic deletion animal models.

**Fig. 4.**
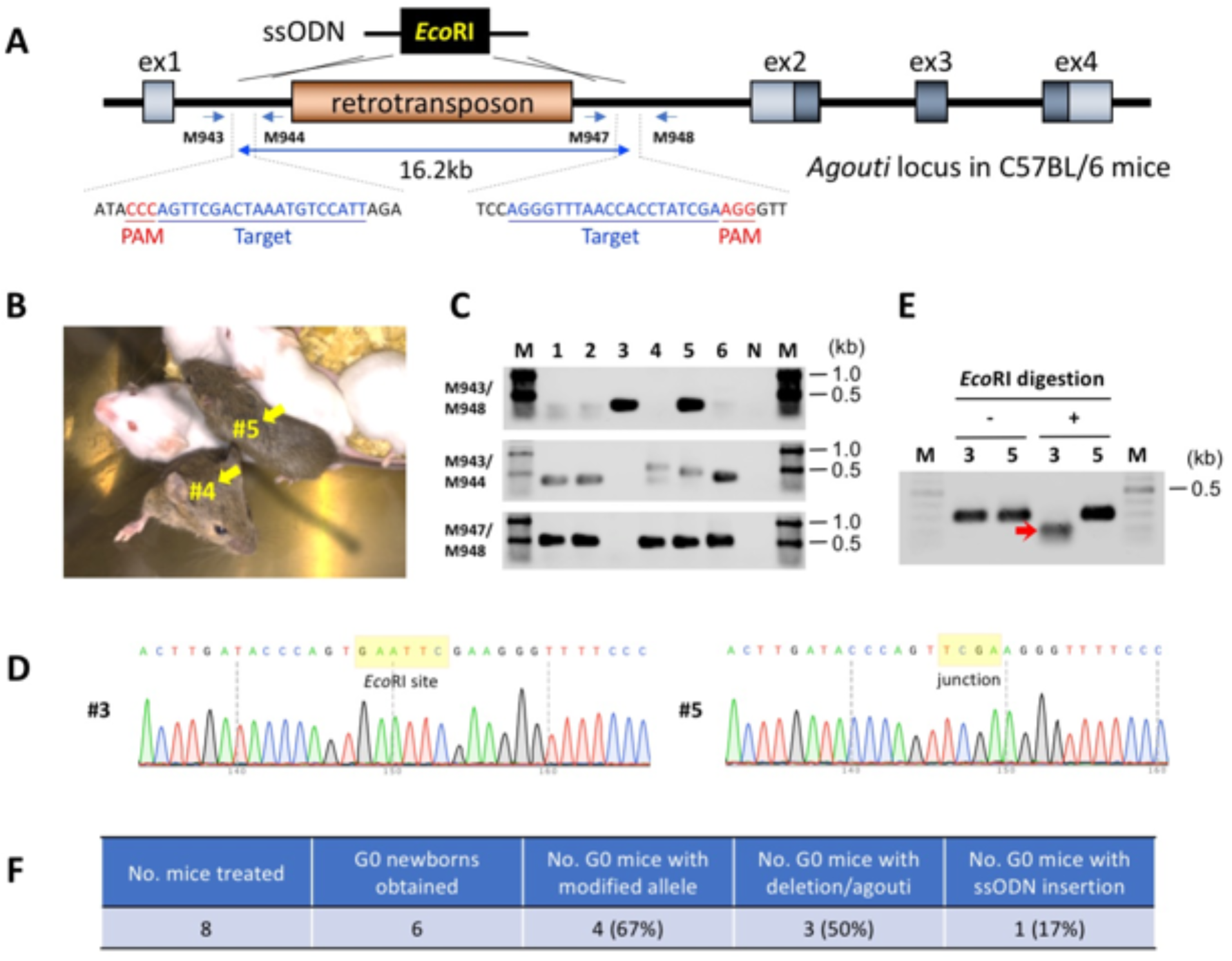
Creating large deletion using the GONAD method. **A.** A schematic diagram showing deletion of 16.2 kb sequence consisting of retrotransposon in the C57BL/6 mouse genome, to restore agouti phenotype. The target sequences and genotyping primers are shown. ssODN containing *Eco*RI site at the middle of the sequence was used. **B.** Representative mice showing rescued-agouti phenotype (indicated by yellow arrows). These mice were recovered through Caesarean section and nursed by ICR foster mother with her own pups. **C.** Genotyping analyses. Expected fragment sizes: M943/M948 = 290- or 295-bp (ssODN knock-in), M943/M944 = 337-bp, M947/M948 = 477-bp. **D.** Direct sequencing results of PCR products amplified from the GO mice. The position of junctional sequences are indicated by yellow rectangles. **E.** *Eco*RI digestion of PCR products amplified from GO mice (G0-#3 and -#5) with the M943/M948 primer set. Red arrow indicates digested fragment. F. Efficiencies of *agouti* gene editing (see **Supplemental Table S4** for detail).

### 6) Knocking-in of long ssDNA donors using *i*-GONAD

We previously demonstrated that knocking-in of longer sequences can be efficiently achieved using ssDNA donors ^18–20^. We tested whether long ssDNA donors can be used with *i*-GONAD to create knock-in alleles. The *Pitx3* and *Tis21* genes were selected for the knock-in experiment to create reporter models with T2A-mCitrine fusion cassettes.

We inserted a 783-bp “T2A-mCitrine” cassette immediately upstream of the stop codon of *Pitx3* gene (**Fig. 5A**). The ssDNA donor was prepared using an *in vitro* Transcription and Reverse Transcription (*iv*TRT) method we previously described ^18^. The concentration of Cas9 protein (1 mg/ml) and crRNA/tracrRNA (30 µM) was the same used in the previous experiment. The ssDNA was used at concentrations of 1.4 or 2.2 ng/µl as donors in the *i*-GONAD procedure. The G0 fetuses were dissected at E12.5 and were observed for fluorescence under a dissecting microscope. The eye-lenses of some fetuses exhibited fluorescence (**Fig. 5B**), suggestive of correct insertion of the fusion cassette. This observation is similar to the previously created knock-in model ^21^. The correct insertion of the cassette at the target site was confirmed by PCR amplification and sequencing of the target region (**Fig. 5C**), which revealed that 15% (5/34) of samples contained the knock-in cassettes (**Fig. 5D** and **5E**). Of note, the other alleles of these five samples contained *indel* mutations. Also, 16 fetuses that did not contain the insertion of the T2A-mCitrine cassette contained *indel* mutations (**Fig. 5E**; **Supplemental Table 5**). Collectively, the percentage of G0 individuals that had been genome-edited (knock-in and/or *indel* mutations) was 62% (21/34).

**Fig. 5.**
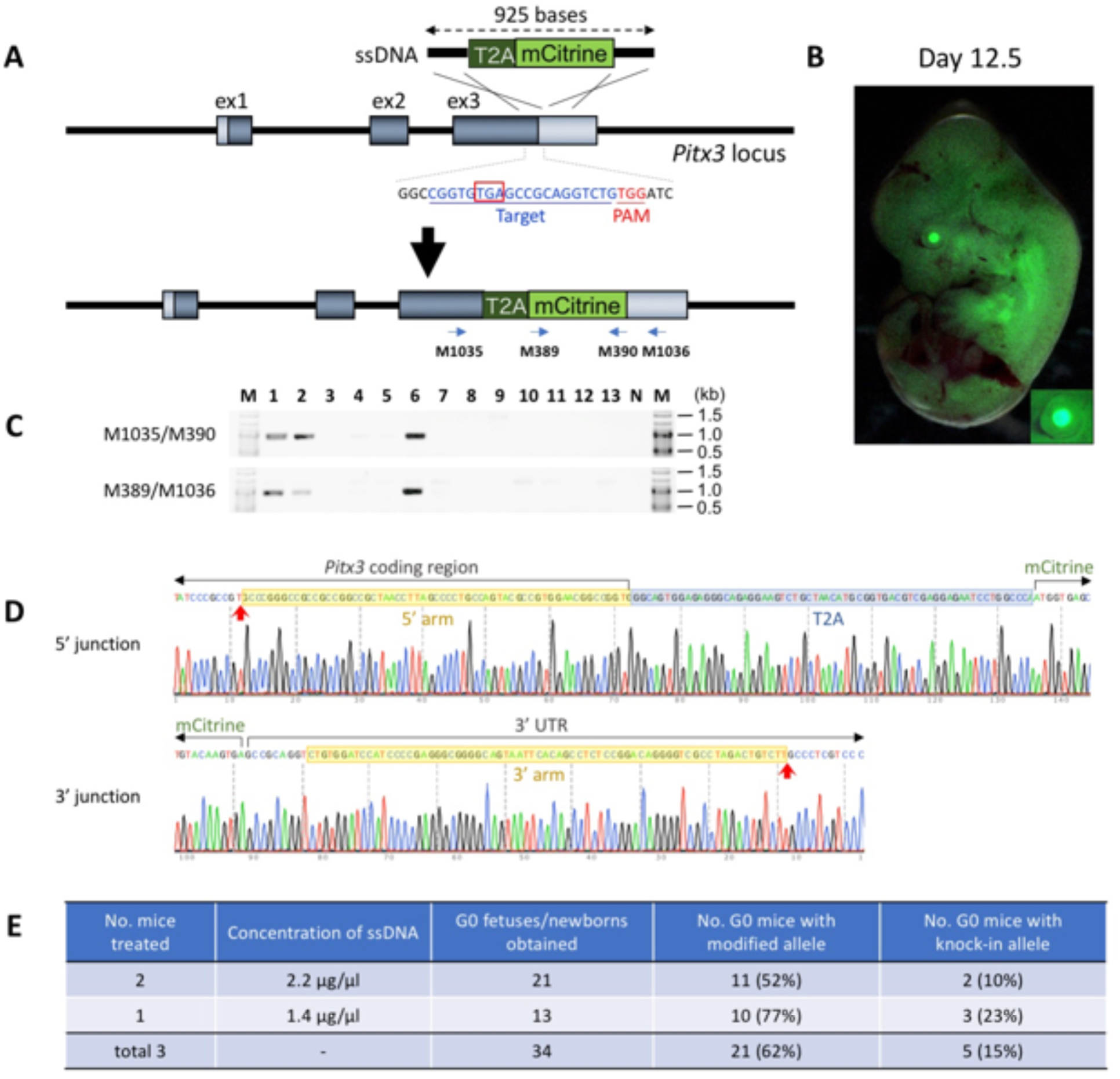
Generation of reporter knock-in mice using the GONAD method, **A.** A schematic diagram showing insertion of “T2A-mCitrine” cassette into *Pitx3* locus. The target sequence and the genotyping primer sets are shown. A 925 bases-long ssDNA synthesized by *iv*TRT method was used as the donor DNA. **B.** mCitrine fluorescence in fetus collected at E12.5. The eye of the fetus is enlarged as an inset. **C.** Example of genotyping analysis of knock-in GO fetuses. Expected fragment sizes: M1035/M390 = 948-bp, M389/M1036 = 956-bp. N: negative control, M: size marker. **D.** A representative sequencing chromatogram showing 5′ and 3′ junctional regions of the inserted cassette. The junctional sequences showing insertion derived from G0-#1 in C is shown. Red arrows indicate junctions between the arms and the genomic sequences. **E.** Genome editing efficiency of the *Pitx3* locus by the GONAD method.

By using the same strategy as knock-in into *Pitx3* gene, we designed a knock-in of a “T2A-mCitrine” cassette to be inserted into *Tis21* gene, immediately upstream of its stop codon (**Supplemental Fig. 3A**). The ssDNA was used at concentrations of 0.85 ng/µl, and G0 fetuses analyzed at E14.5. 14 fetuses were recovered and observed for fluorescence and genotyped by PCR. One of the fetuses exhibited fluorescence in developing CNS (**Supplemental Fig. 3B**) similar to a *Tis21* knock-in model created previously ^22^ and this fetus had the correct knock-in sequence as analyzed by PCR and sequencing. Two other fetuses that did not show fluorescence contained partial insertion of the knock-in cassette (**Supplemental Fig. 3C** to **3E**) and contained *indel* mutations in their second alleles (**Supplemental Fig. 3E**; **Supplemental Table 5**). One more fetus that did not show knock-in had an *indel* mutation. Taken together, the percentage of G0 pups that were genome-edited was 29% (4/14: 1/14 knock-in and 3/14 *indel* mutations).

### 7) GONAD using Cpf1 protein

Recently, Cpf1 derived from *Acidaminococcus sp.* has been added as an genome editing tools and used for creating genome edited animal models ^23,24^. We tested if Cpf1 protein can be used in GONAD, which was performed by instilling 6.3 µM of Cpf1 protein into the oviductal lumen of two pregnant ICR mice together with 30 µM of gRNA targeting the *Hprt* locus. A total of 11 embryos were isolated at E13.5, and the presence or absence of *indel* was analyzed by PCR and sequencing. One fetus did not amplify PCR fragment probably due to deletion of primer biding site(s), and surprisingly, all the other fetuses contained *indel* mutation. The results show that all of the G0 offspring recovered after the *i*-GONAD procedure (100% [11/11]) were genome-edited, indicating that the *i*-GONAD can yield very high efficiencies of genome editing even when Cpf1 is applied (**Supplemental Table 6**).

### 8) The animals used for GONAD procedure retain reproductive function

GONAD method does not require euthanasia of pregnant female mice, unlike in the traditional approaches where females need to be sacrificed for isolating zygotes for introduction of genome editing components *ex vivo.* We investigated if female mice subjected to GONAD retain their reproductive functions. Three female mice that underwent GONAD and delivered genome-edited pups (mice-#3, -#5, and -#11 in **Supplemental Table 2**) were naturally mated to fertile male mice. Two of these (67%; mice-#3 and -#11) became pregnant and successfully delivered 9 and 12 pups, respectively (**Supplemental Fig. 4**). These data suggest that oviducts in these mice retained normal reproductive function, allowing fertilization and subsequent tubal transport of fertilized eggs to uteri.

## Discussion

We previously demonstrated a proof of principle genome editing method called GONAD. GONAD can be performed on zygotes *in situ* and thus bypasses the steps of isolation of zygotes, their *ex vivo* handling, and their subsequent transfer to recipient females, which are the inevitable steps of animal genome editing that were developed and have been practiced for over three decades. In this study, we made several improvements to the GONAD method, making it highly suitable for routine creation of genome edited animal models. First, we assessed the optimal time of pregnancy and show that Day 0.7 would be better for reducing mosaicism because the embryos are still at one cell stage. Second, replacing Cas9 mRNA with Cas9 protein and replacing sgRNA with separated guides (crRNA + tracrRNA) in the approach, termed *i*-GONAD, enhanced genome editing efficiency that is comparable, or even better than, the microinjection-based approach. Third, the *i*-GONAD can be used for creating large deletion, point mutation- and larger cassettes-knock-in animal models. Fourth, the females used for *i*-GONAD retain fully functional reproductive capability. Fifth, the Cpf1 nuclease can also be used in the *i*-GONAD method. Lastly, we demonstrate that *i*-GONAD can be performed using different types of commercially available electroporators. Because the electroporaters are nearly ten times less expensive than microinjection set ups, and do not require specialized personnel to operate them, the *i*-GONAD method can be readily adapted at many laboratories that lack one or both.

The GONAD on Day 0.7 (corresponding to late one-cell stage) was shown to be effective for genome editing. There are several advantages of performing GONAD at Day 0.7. First, it results in reduced mosaicism for the mutated target gene (**Fig. 2B**), when compared to that on Day 1.5 (corresponding to 2-cell stage) ^10^. The zygotes at Day 0.7 will be surrounded by a fewer number of cumulus cells (**Fig. 1A**), and thus are easily accessible for delivery of genome editing components (CRISPR-related nucleic acids/protein). We saw that our experiments at Day 0.4 did not elicit effective delivery of components via electroporation, probably because zygotes are surrounded by a cluster of cumulous cells at this stage of pregnancy. Second, the oviduct on Day 0.7 has a distinctly visible ampulla in many cases, which will facilitate micropipette-aided instillation of solutions while observing under a dissecting microscope. Third, in comparison to our previously reported GONAD procedure, the time point of Day 1.5, the amount of a solution instilled at Day 0.7 can be reduced to 1.0~1.5 µl per oviduct; the amount of liquid needed to fill the oviduct of Day 1.5 pregnant female is often more than 1.5 µl of solution ^10,11^. Forth, we speculate that because the zygotes at Day 0.7 would be in a smaller compacted space in the ampulla of the oviduct, the genome editing components may leach to zygotes more effectively than at Day 1.5.

Accumulated data from zygote-injection or *ex vivo* electroporation-based genome editing have established that RNP elicits superior genome editing efficiencies than mRNA/sgRNA ^7,14^. GONAD performed using RNP components in this study also achieved up to ~100% of genome editing efficiency, whereas the efficiencies reached only up to 31% when using sgRNA/Cas9 mRNA components (**Fig. 2F**). Another advantage of the RNP platform is that all components can be custom-made, which will reduce variations resulting in reagents when prepared in individual labs. The commercial reagents can also be received as lyophilized form for RNA components and Cas9 protein can be purchased at higher concentration; these features allow preparation of electroporation mixes at any desired concentrations. For GONAD procedures, the concentration of reagents required in the electroporation mix are typically much higher than the mixes used for direct zygote injection, because large quantities of electroporation mix are spread out within and around the ampulla, so that a concentration of the solution high enough to cause editing will likely be in contact with most free-floating zygotes. The fact that 100% of zygotes were genome edited in some females indicate such a possibility.

The knock-in efficiency of ssODN donors in the *i*-GONAD-treated samples was ~50% (**Fig. 3E**). By using the same combination of locus and genetic modification, we directly compared this efficiency with the microinjection genome editing method, and the efficiency was 33% (**Supplemental Table 3**). The number of animals needed for *i*-GONAD are three times less than those needed for microinjection-based approaches, at least when using an ICR strain. More loci must be tested to assess the comparable efficiency of the two methods. We also successfully inserted a long donor fragment into the targeted locus using the *i*-GONAD method. Long ssDNA donors were prepared using the *iv*TRT method described in our highly efficient knock-in method, “*Easi*-CRISPR” ^18–20^. Since a large amount of ssDNA is required for the *i*-GONAD experiment, we used the spin column-based nucleic acid purification instead of gel purification where loss of sample recovery is very high. Since the microinjection approach does not require higher concentration, gel purification is typically followed for zygote-microinjection experiments ^18^. The column purified ssDNA (922~925 bases) exhibited a single-band after separation using agarose gel electrophoresis, which produced knock-in mice when used as the donor in the GONAD procedure (**Fig. 5**). Column purified ssDNAs can also be used as donors for creating floxed mice (i.e., mice with a gene locus flanked by *loxP* using *Easi*-CRISPR). Unlike ssODN knock-in, the efficiency of inserting a long donor fragment with the *i*-GONAD method appeared to be low (up to 15%) compared to microinjection (25~67%) ^20^. However, *i*-GONAD would be considered superior to microinjection because it uses far fewer animals because it does not require a second set of animals (vasectomized males and pseudopregnant females) to carry the manipulated zygotes to full term. This second step is inevitably required using the microinjection or *ex vivo* electroporation approaches. Thus, the *i*-GONAD method can be used as an alternative to zygote microinjection for creating knock-in alleles. *i*-GONAD is also a convenient and simple method, as stated above, for production of genome-edited animals.

In this study, we showed that *i*-GONAD can be used to rescue pigmentation defects in albino mice (ICR strain) and black mice (C57BL/6 strain) by correction of point mutation in the *Tyr* gene and elimination of retrotransposon sequence in the *agouti* gene, respectively. Such genetic alterations are quite common in many human genetic disease ^25,26^ and our strategy can be applicable to human germline gene therapy to correct disease causing mutations. Insertion of long sequences will be also useful as one of the gene therapy strategies based on the addition of a functional gene ^27^. Considering that human germline gene therapy will often be coupled with *ex vivo* handling of embryos, including an *in vitro* cell culture step that could cause epigenetic changes of gene expression and affect fetal development ^28,29^, *i*-GONAD, which does not require *ex vivo* handling or sacrifice of GONAD-treated females, can offer a highly promising approach as a human germline gene therapy tool in the future.

To date, several reports regarding production of genome-edited rodents through an *in vitro* electroporation system have been provided ^4–9^. The GONAD method is a step beyond these, given that it directly and very smoothly delivers genome editing nucleic acids and CRISPR components into embryos *in situ*. The GONAD method offers even more advantages over *in vitro* electroporation-based genome editing methods, as we have demonstrated: 1) GONAD does not require *ex vivo* handling of embryos, including *in vitro* cultivation of isolated embryos, pseudopregnant female mice that allow implantation of *ex vivo*-treated embryos, or vasectomized males to produce pseudopregnant females. This feature would be advantageous if the animal of interest is sensitive for *ex vivo* handling and/or difficult to prepare surrogate mothers. 2) GONAD-treated females need not be sacrificed for zygote isolation. Furthermore, we found that GONAD-treated females retain the reproductive functions and can become pregnant after delivering pups from the GONAD procedure, suggesting the possibility that the females can be re-used for a second GONAD procedure. This is a very important feature in the situations such as: (i) the animals used for GONAD experiments are valuable, and; (ii) another genetic manipulation can be performed immediately in a newly developed mouse line. This avoids the laborious requirement of expanding the line to produce hundreds of zygotes for performing the second genetic change when using microinjection or *ex vivo* electroporation approaches.

## Methods

### CRISPR reagents

CRISPR guide RNAs were designed using CRISPR.mit.edu or CHOPCHOP (**Supplemental Table 7**). The sgRNA for *Foxe3* was synthesized as described previously ^10^ using the primer sets (M1055/M939) and the pUC57-sgRNA vector as template (Addgene plasmid number: #51132). The mRNAs for eGFP and Cas9 were *in vitro* transcribed as previously described ^10,11^. The synthetic crRNA and tracrRNA were commercially obtained as Alt-R^TM^ CRISPR guide RNAs from Integrated DNA Technologies [IDT]) together with Cas9 protein (Alt-R^TM^ S.p. Cas9 Nuclease 3NLS). The ssODN donors (for *Tyr* rescue experiment [*Tyr*-rescue] and *agouti* rescue experiment [*agouti*-rescue]) were custom synthesized from IDT. Long ssDNA donors (for *Pitx3* and *Tis21* reporters) were prepared from the dsDNA templates using the *iv*TRT method described previously ^18^ with slight modifications. The “T2A-mCitrine cassette” was amplified from original vector (pP200) with primer sets (M1051/M1052 for *Pitx3* and M1053/M1054 for *Tis21*) and inserted into the *Sma*I site of pUC119, resulting in pP206 (for *Pitx3*) and pP209 (for *Tis21*). The templates for RNA synthesis were amplified from these vectors with primer sets (PP226/M272 for *Pitx3* and PP227/M272 for *Tis21*) and RNAs were synthesized using T7 RiboMax Express Large Scale RNA Production System (Promega). The RNAs were purified using MEGAclear Kit (Ambion) and the cDNAs were generated using SuperScript III Reverse Transcriptase (for *Pitx3*) or SuperScript IV Reverse Transcriptase (for *Tis21*; Life Technologies) with the primer (PP226 for *Pitx3* and PP227 for *Tis21*). The final step of gel extraction, as done for purifying cDNA for microinjection, was excluded in order to obtain sufficiently higher concentration of the final ssDNA. Instead, spin column-based nucleic acid purification using NucleoSpin Gel and PCR Clean-up (MACHEREYNAGEL) was performed. After ethanol precipitation, the DNA pellet was dissolved in EmbryoMax Injection Buffer (Millipore). The sequences for primers and ssODNs are shown in **Supplemental Table 8**.

### Mice

All mice were maintained at the Tokai University School of Medicine animal facility. Adult ICR and C57BL/6J mice were obtained from CLEA Japan, Inc. (Tokyo, Japan). All the animal experiments were performed in accordance with institutional guidelines and were approved by The Institutional Animal Care and Use Committee at Tokai University (Permit Number: #154014, #165009).

### Preparation of CRISPR electroporation solutions

The solution contained *in vitro*-synthesized RNAs or purchased crRNA/tracrRNA and Cas9 protein and/or ssODN/ssDNA were prepared. Cas9 mRNA/sgRNA mixture was prepared as we previously described ^10^. 0.05% of trypan blue (Nacalai tesque Inc., Kyoto, Japan) used as a marker for successful instillation, was used only when eGFP mRNA was used or Cas9 was supplied as mRNA. Lyophilized ssODNs (Ultremer; commercially procured from IDT) were re-suspended in nuclease-free water to a concentration of 10 µg/µl. Lyophilized crRNA and tracrRNA were first re-suspended in RNase-free Duplex Buffer to a concentration of 200 µM. Equal volume of crRNA and tracrRNA were combined in a 1.5 ml tube and were heated in a thermocycler to 94 °C for 2 min and then placed at room temperature for about 10 min. The annealed crRNA and tracrRNA were mixed with Cas9 protein and/or ssODN/ssDNA so that the final concentrations of components were 30 µM (for crRNA/tracrRNA), 1 mg/ml (for Cas9 protein), 2 µg/µl (for ssODN), and 0.85~2.2 µg/μl (for ssDNA). Cpf1 crRNA (MmHPRT-273-S: 5'-GTGCCCTCTTCTGGCCTGCCA-3') was a kind gift from IDT. Lyophilized crRNAs were first re-suspended in RNase-free water to a concentration of 100 µM, and then heated in a thermocycler to 95 °C for 5 min and placed at room temperature for about 10 min. Cpf1 protein (IDT) was mixed with crRNA so that the final concentrations of components were 30 µM (for crRNA) and 6.3 µM (for Cpf1 protein). The electroporation solution was occasionally diluted using Opti-MEM (Thermo Fisher Scientific) to adjust the volume to 1.5 µl/oviduct.

### GONAD procedure

The super-ovulated C57BL/6J female mice were mated to adult male C57BL/6J, or estrous ICR female mice were mated to adult male ICR mice without super-ovulation. Presence of copulation plugs was confirmed by visual inspection next morning and the females having plugs were designated as day 0.5 of gestation at 12:00 and day 0.7 of gestation at 16:00, and used for the electroporation experiments.

Surgical procedures were performed on anesthetized females at Day 0.7 of pregnancy (corresponding to late 1-cell stage zygotes; at 16:00 of the same day when plugs were confirmed) under observation using a dissecting microscope (SZ11; Olympus, Tokyo, Japan), as described previously ^10,11^ with slight modification. The ovary/oviduct/uterus were exposed after making an incision at the dorsal skin. Approximately 1.0-1.5 µL of electroporation solution (pre-warmed at 37°C for 10min) was injected into the oviductal lumen from upstream of the ampulla using a micropipette. The micropipette apparatus consisted of a glass capillary needle (pulled using an electric puller: P-97/IVF; Sutter Instrument Co., Novato, CA, USA) and a mouthpiece attached to the needle. Immediately after the injection of solution, the oviductal regions were covered with a piece of wet paper (Kim wipe; Jujo-Kimberly Co. Ltd., Tokyo, Japan) soaked in phosphate-buffered saline (PBS), and were grasped in tweezer-type electrodes (CUY652-3 [NEPA GENE Co. Ltd., Ichikawa, Chiba, Japan] for T820 and NEPA21, and LF650P3 [BEX Co. Ltd., Tokyo, Japan] for CUY21EDIT II). The electroporation was performed using a square-wave pulse generator T820 (BTX Genetronics Inc.), or NEPA21 (NEPA GENE), or CUY21EDIT II (BEX). The electroporation parameters were as follows; eight square-wave pulses with a pulse duration of 5 msec, pulse interval of 1 sec and an electric field intensity of 50 V for T820, Poring pulse; 50 V, 5 msec pulse, 50 msec pulse interval, 3 pulse, 10% decay (± pulse orientation) and Transfer pulse; 10 V, 50 msec pulse, 50 msec pulse interval, 3 pulse, 40% decay (± pulse orientation) for NEPA21, and Pd V: 60V, Pd A: 200mA, Pd on: 5.00ms, Pd off: 50ms, Pd N: 3, Decay: 10%, DecayType: Log for CUY21EDIT II. After the electroporation, the oviducts were returned to their original position and the incisions were sutured. The animals were monitored for anesthesia recovery and were housed for further analysis.

### Microinjection

CRISPR components were mixed in EmbryoMax Injection Buffer. Final concentrations of Cas9 protein, crRNA/tracrRNA, and ssODN (for *Tyr* rescue) were 50 ng/µl, 0.61 µM, and 10 ng/µl, respectively. Unfertilized oocytes isolated from super-ovulated female mice (ICR) were subjected to *in vitro* fertilization (IVF) with spermatozoa freshly isolated from an ICR male mouse. Microinjection of the mixture was performed into pronuclei of *in vitro* fertilized eggs. The injected embryos were transferred into the oviduct of pseudopregnant ICR females to allow further development. The resulting fetuses (Day 13.5) were recovered and subjected to genotyping analysis.

### Observation of mCitrine fluorescence

The fetuses recovered were observed using a fluorescence stereomicroscope with filter for GFP (Olympus SZX7 with SZX-MGFPA) for detecting the mCitrine fluorescence.

### Analysis of CRISPR/Cas9-induced mutations and insertions

Genomic DNAs were isolated from the midgestational fetuses or the ear-piece of live bron mice using All-In-One Mouse Tail Lysis Buffer (ABP-PPMT01500; KURABO, Osaka, Japan) through incubation at 55°C for 3 h or overnight and subsequent inactivation at 85°C for 45 min. The PCR for amplification of target loci *Foxe3*, *Tyr*, *agouti*, and *Hprt* was performed in a total of 10 µl solution containing 5 µl of 2 x GC buffer I, 0.2 mM dNTP, 1 µl of the crude lysate, the primer pairs (**Supplemental Table 8**), and 0.125 U of TaKaRa r-Taq (TaKaRa) using denaturation (95°C for 5 min), 35 cycles of 95°C for 45 sec, 58°C for 30 sec and 72°C for 1 min, and extension (72°C for 5 min). For amplification of target loci *Pitx3* and *Tis21*, PCR amplifications were performed using PrimeSTAR HS DNA Polymerase (TaKaRa) in a total of 10 µl solution containing 2 µl of 5×PrimeSTAR buffer I, 0.2 mM dNTP, 1 µl of the crude lysate, the primer pairs (**Supplemental Table 8**), and 0.25 U of PrimeSTAR HS DNA Polymerase using denaturation (94°C for 3 min), 35 cycles of 98°C for 10 sec, 62°C for 5 sec and 72°C for 2 min, and extension (72°C for 10 min). Direct sequencing was performed using the PCR products and the primers listed in **Supplemental Table 8**.

## Acknowledgements

This work was supported in part by Grant-in-Aid for Young Scientists (B) (16K18821) from Japan Society for the Promotion of Science (JSPS) to H.M. MO acknowledges the funding support by the 2014 Tokai University School of Medicine Research Aid, MEXT-Supported Program for the Strategic Research Foundation at Private Universities 2015-2019 (PI: Y.Inagaki), Research and Study Project of Tokai University General Research Organization, 2016-2017 Tokai University School of Medicine Project Research, and Grant-in-Aid for challenging Exploratory Research (15K14371) from JSPS. This work was also supported by Grant-in-Aid for Scientific Research (B) (16H05049) from JSPS to S.N. We thank G. Takahashi (Tokyo University of Agriculture) for designing of gRNA and primer set for *Foxe3* gene. The authors gratefully acknowledge the contribution of the staff of Support Center for Medical Research and Education, Tokai University for microinjection and prepareing pregnant mice (A. Nakamura, S. Ogiwara, and Y. Ishikawa), and for sequencing. Cpf1 protein and the guide RNAs were gifts from IDT.

## Authors' contributions

M.O, S.N, K.W, C.B.G and M.S conceived this study, M.O, K.W and M.S designed the experiments, M.O, H.M, N.A and M.S performed experiments. M.O, M.S and C.B.G wrote the manuscript with input from other authors.

